# Functional properties of the spike glycoprotein of the emerging SARS-CoV-2 variant B.1.1.529

**DOI:** 10.1101/2021.12.27.474288

**Authors:** Qian Wang, Saumya Anang, Sho Iketani, Yicheng Guo, Lihong Liu, David D. Ho, Joseph G. Sodroski

## Abstract

The recently emerged B.1.1.529 (Omicron) SARS-CoV-2 variant has a highly divergent spike (S) glycoprotein. We compared the functional properties of B.1.1.529 S with those of previous globally prevalent SARS-CoV-2 variants, D614G and B.1.617.2. Relative to these variants, B.1.1.529 S exhibits decreased processing, resulting in less efficient syncytium formation and lower S incorporation into virus particles. Nonetheless, B.1.1.529 S supports virus infection equivalently. B.1.1.529 and B.1.617.2 S glycoproteins bind ACE2 with higher affinity than D614G S. The unliganded B.1.1.529 S trimer is less stable at low temperatures than the other SARS-CoV-2 spikes, a property related to spike conformation. Upon ACE2 binding, the B.1.1.529 S trimer sheds S1 at 37 degrees but not at 0 degrees C. B.1.1.529 pseudoviruses are relatively resistant to neutralization by sera from convalescent COVID-19 patients and vaccinees. These properties of the B.1.1.529 spike glycoprotein likely influence the transmission, cytopathic effects and immune evasion of this emerging variant.

## INTRODUCTION

The continuing coronavirus disease 2019 (COVID-19) pandemic has stimulated the implementation of a number of countermeasures, including vaccines, therapeutic antibodies and antiviral agents. Vaccines that elicit neutralizing antibodies against the spike (S) glycoprotein of the etiologic agent of COVID-19, severe acute respiratory syndrome coronavirus 2 (SARS-CoV-2), have effectively reduced the probability of infection and death from this virus (Baden et al., 2021; Dai and Gao, 2021; Polack et al., 2020; Sadoff et al., 2021). However, the emergence of a new SARS-CoV-2 variant in Botswana and South Africa in mid-November 2021, coupled with its rapid spread throughout the world, has raised concerns (Grabowski et al., 2021; Pulliam et al., 2021; Scott et al., 2021). The high rate of transmission has caused the World Health Organization to classify this variant as a Variant of Concern (VOC), and it has been termed Omicron (formally, B.1.1.529), now found in over 70 countries (WHO, 2021).

B.1.1.529 was found to have an unprecedented level of divergence, with more than 55 mutations compared to the ancestral Wuhan-Hu-1 strain. Of particular importance to antibody-mediated protection, more than 30 of these mutations alter the sequence of viral spike (S) glycoprotein (Scott et al., 2021). Several of these changes in S, such as K417N and N501Y, have been previously demonstrated to confer resistance to antibody neutralization in other SARS-CoV-2 variants (Wang et al., 2021a; Wang et al., 2021b). Residue Glu 484, which is altered to Lys in the B.1.351 and P.1 variants, is an Ala in B.1.1.529, resulting in a similar loss of sensitivity to a subset of antibodies (Liu et al., 2021). These features suggest that the B.1.1.529 virus may be highly adapted to resist antibodies directed against the ancestral SARS-CoV-2.

Recent reports have confirmed this prediction. Several *in vitro* studies have demonstrated that B.1.1.529 evades several monoclonal antibodies, including some used clinically, and is less effectively neutralized by antibodies elicited by SARS-CoV-2 infection and vaccines (Doria-Rose et al., 2021; Garcia-Beltran et al., 2021; Gruell et al., 2021; Hoffmann et al., 2021; Liu et al., 2021; Planas et al., 2021). The degree of B.1.1.529 resistance to antibodies seems to be greater than that observed in any variant to date, and consequently many current vaccinees may be at risk of breakthrough infections or disease (Cong et al., 2021; Lu et al., 2021). Data from the United Kingdom and South Africa indicate a substantial decline in the efficacy of the BNT162b2 vaccine (Pfizer), contextualizing the implications of the experimentally measured resistance (Abu-Raddad et al., 2021; Madhi et al., 2021; Sadoff et al., 2021).

Given the altered epidemiology of B.1.1.529 relative to other SARS-CoV-2 variants, we investigated the functional properties of the S glycoprotein of B.1.1.529. By comparison to the current predominant strain, B.1.617.2 (Delta variant), and its ancestor D614G (Wuhan-Hu-1 strain with a D614G change in S), we find that the alterations in B.1.1.529 confer unique characteristics on the spike. Notably, B.1.1.529 has significantly reduced processing of its S glycoprotein into S1 and S2 subunits, which decreases syncytium formation, a major cytopathic effect of SARS-CoV-2 infection. Another apparent consequence of poor S processing is a reduced incorporation of spike into pseudovirus particles, compared with that of D614G and B.1.617.2. Like other viruses descended from D614G, B.1.1.529 exhibits a higher affinity of its spike for the cognate receptor ACE2 and efficiently mediates pseudovirus infection of ACE2-expressing cells. The B.1.1.529 spike resists soluble ACE2-induced shedding of the S1 exterior glycoprotein at lower temperatures. After extended incubation on ice, the B.1.1.529 spike glycoprotein spontaneously sheds the S1 subunit and loses its ability to support virus entry. The spontaneous shedding of S1 and the cold inactivation of the virus spike are related to the conformation of the receptor-binding domains (RBDs) on the spike glycoprotein. We confirm that the B.1.1.529 S glycoprotein is less sensitive to neutralization by sera from convalescent COVID-19 patients and recipients of two doses of the mRNA-1273 vaccine (Moderna). The binding of the sera to recombinant S trimers indicates that many antibodies in the convalescent patient sera bind conserved epitopes on the B.1.1.529 spike that are not available as neutralization targets. Serum binding to the D614G and B.1.1.529 RBDs correlated better with the virus neutralization activity of the sera. Collectively, our study reveals biological properties of the divergent B.1.1.529 spike that may contribute to its rapid transmission, altered pathogenesis and immune escape.

## RESULTS

### S glycoprotein expression, post-translational processing and function

The significant number of changes in the B.1.1.529 spike glycoprotein (**Figure 1A**) led us to compare its properties with those of spike glycoproteins from two other globally prevalent SARS-CoV-2 strains, D614G and B.1.617.2. We studied the expression and processing of the D614G, B.1.617.2 and B.1.1.529 S glycoproteins. We generated vesicular stomatitis virus (VSV) and lentivirus (HIV-1)-based pseudovirus particles as previously described, and examined the incorporated S glycoproteins by Western Blot (Schmidt et al., 2020; Wang et al., 2021c). In both pseudoviral systems, relative to the D614G spike, the B.1.617.2 S glycoprotein was proteolytically processed more efficiently, whereas the B.1.1.529 S glycoprotein exhibited significantly less cleavage (**Figure 1B and C**). For all these SARS-CoV-2 S glycoproteins, the cleaved (S1 and S2) glycoproteins were preferentially incorporated into pseudovirus particles. Furthermore, the decrease in B.1.1.529 S glycoprotein cleavage was associated with reduced incorporation of spike glycoproteins into pseudovirus particles (**Figure 1B**).

**Figure 1.**
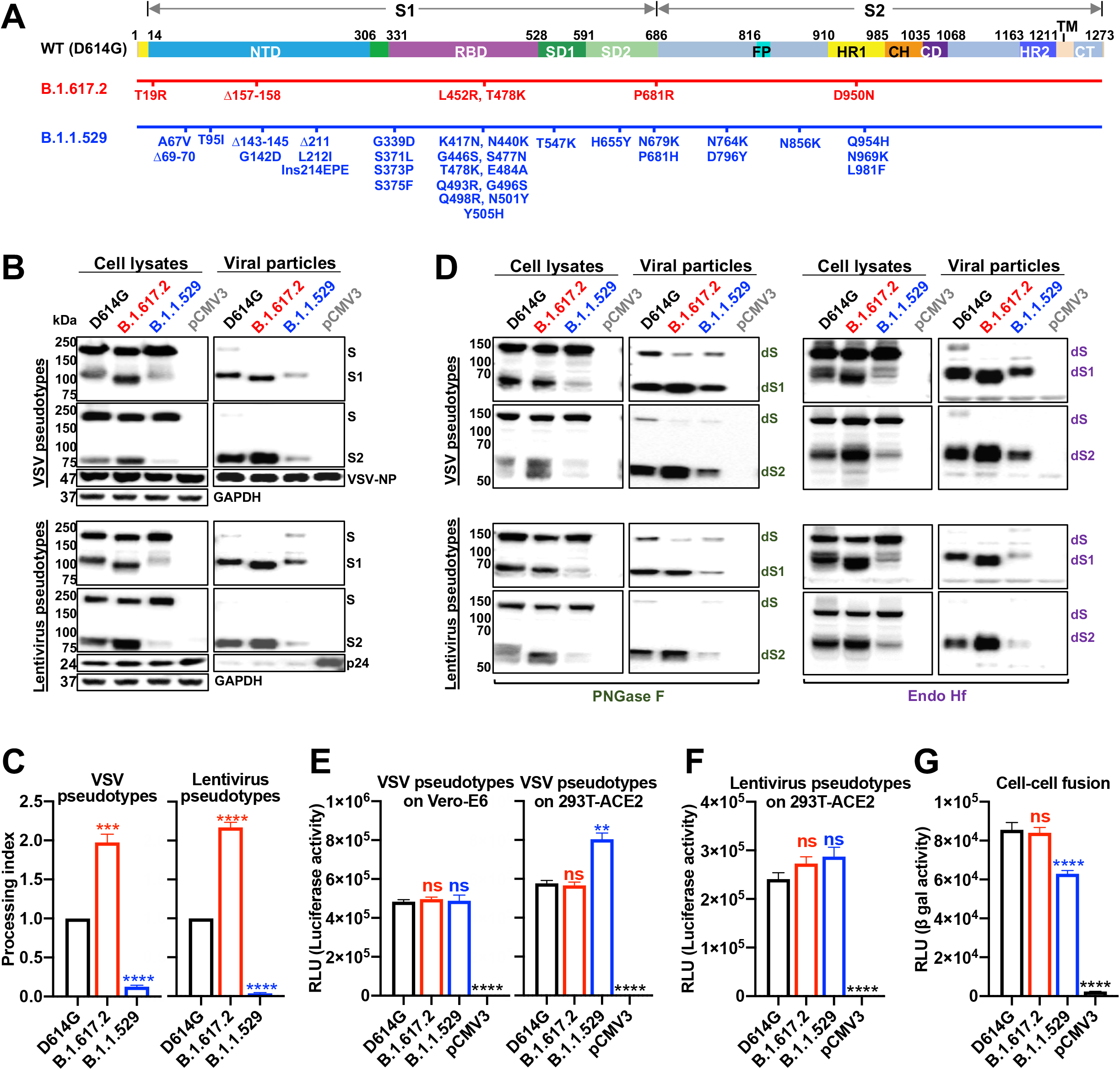
SARS-CoV-2 variant S glycoproteins expression, post-translational processing and function. **(A)** A schematic representation of the wild-type (D614G) S glycoprotein is shown in the upper figure. The NTD, RBD, and subdomains 1 and 2 (SD1 and SD2) of the S1 glycoprotein are indicated. The fusion peptide (FP), heptad repeat regions (HR1 and HR2), central helical region (CH), connector domain (CD), transmembrane region (TM), and cytoplasmic tail (CT) of the S2 glycoprotein are indicated. The amino acid changes in the B.1.617.2 and B.1.1.529 spikes, relative to the D614G S glycoprotein, are shown in the diagrams beneath the schematic. **(B)** HEK293T cells producing VSV (upper panels) and lentiviral (lower panels) particles pseudotyped with the indicated SARS-CoV-2 S glycoproteins were used to prepare cell lysates (left panels) and viral particles (right panels), which were Western blotted with rabbit anti-S1 and anti-S2 antibodies. The samples were also Western blotted with anti-VSV NP (upper panels), anti-HIV p17/p24/p55 (lower panels), and anti-GAPDH antibodies. The Western blots shown are representative of those obtained in three independent experiments. Cells transfected with the pCMV3 vector were used as a negative control. **(C)** Processing indices were calculated from Western blots of cell lysates from three experiments. **(D)** The cell lysates and viral particle preparations from **(B)** were treated with PNGase F (left panel) and Endo Hf (right panel), and the deglycosylated samples were Western blotted with rabbit anti-S1 and anti-S2 antibodies. Cells transfected with the pCMV3 vector were used as a negative control. The deglycosylated forms of the S glycoprotein precursor (dS), S1 subunit (dS1), and S2 subunit (dS2) are indicated. The Western blots shown are representative of those obtained in two independent experiments. **(E)** The infectivity of VSV vectors pseudotyped by the indicated SARS-CoV-2 S glycoproteins was determined on Vero-E6 cells (left panel) and 293T-ACE2 cells (right panel). The pCMV3-transfected cells serve as a negative control. The results shown are representative of those obtained in three independent experiments. **See also figure S1**. **(F)** The infectivity of lentivirus vectors pseudotyped by the indicated SARS-CoV-2 S glycoproteins was determined on 293T-ACE2 cells. The pCMV3-transfected cells serve as a negative control. The results shown are representative of those obtained in two independent experiments. Data are presented as mean ± SEM. For statistical analysis, a Student’s unpaired *t* test was used to compare the values to those obtained for the wild-type D614G S glycoprotein (**p < 0.01; ***p < 0.001; ****p < 0.0001; ns - not significant).

Next, we examined the glycosylation of these three SARS-CoV-2 S glycoproteins in the two pseudovirus systems. Cell lysates and pseudovirus particles were treated with either PNGase F or Endo Hf, and then visualized by Western blotting (**Figure 1D**). The S glycosylation patterns of the VSV and lentivirus pseudotypes were similar. As previously observed for SARS-CoV-2 spike glycoproteins (Nguyen et al., 2020; Wang et al., 2021c; Zhang et al., 2021), the vast majority of the uncleaved S glycoprotein in cell lysates contained only high-mannose and hybrid glycans, whereas the S1 and S2 glycoproteins on pseudovirus particles were highly modified by complex carbohydrates. Following PNGase F digestion, the S2 glycoproteins incorporated into pseudovirus particles were more homogeneous than those in cell lysates; post-translational modifications of S2 other than N- or O-linked glycosylation account for this difference (Zhang et al., 2021). The untreated and the Endo Hf-treated S1 from B.1.617.2 migrated slightly faster than those of D614G and B.1.1.529, which may be due to a missing N-linked glycosylation site resulting from the T19R change in the N-terminal domain.

To determine if the reduced cleavage and virion incorporation of the B.1.1.529 spike affected viral entry, we quantified the infectivity of the pseudoviruses on Vero-E6 and 293T-ACE2 target cells. For the VSV pseudotypes, no differences among the D614G, B.1.617.2 and B.1.1.529 spikes were observed for infection of Vero-E6 target cells, and the B.1.1.529 S pseudotypes exhibited slightly enhanced infectivity with 293T-ACE2 target cells (**Figure 1E**). The B.1.1.529 pseudoviruses infected 293T cells transiently expressing different levels of human ACE2 comparably to the D614G and B. 1.617.2 pseudoviruses **(Figure S1)**. No significant differences were observed in the infectivity of lentiviruses pseudotyped by the three SARS-CoV-2 S glycoproteins for 293T-ACE2 cells (**Figure 1F**). Collectively, although the B.1.1.529 S glycoprotein exhibits reduced processing and decreased spike incorporation into pseudovirus particles, its ability to mediate virus infection is not compromised.

To evaluate the efficiency of cell-cell fusion mediated by the D614G, B.1.617.2 and B.1.1.529 S glycoproteins, we cocultivated COS-1 cells expressing the S glycoproteins and alpha-gal with 293T cells expressing human ACE2 and omega-gal for 4 hours. The formation of syncytia between the S-expressing cells and ACE2-expressing cells results in the activation of β-galactosidase. Cell-cell fusion mediated by the B.1.1.529 S glycoprotein was lower than the levels observed for the D614G and B.1.617.2 S glycoproteins **(Figure 1G)**. This result is consistent with the expectation that the process of syncytium formation is mediated by cleaved S1/S2 glycoprotein trimers on the surface of the expressing cell (Nguyen et al., 2020). The decreased processing of the B.1.1.529 S glycoprotein results in lower levels of mature S1/S2 trimers in expressing cells, compared with the levels of the D614G and B.1.617.2 S glycoproteins, likely accounting for the observed reduction in cell-cell fusion.

### ACE2 binding and soluble ACE2-induced S1 shedding of variant spikes

We investigated the interaction of the D614G, B.1.617.2 and B.1.1529 S variants with ACE2. We tested the neutralization of D614G, B. 1.617.2, and B.1.1.529 pseudotypes by soluble ACE2 (huACE2-Fc), and observed enhanced neutralization of B.1.617.2 and B.1.1.529 pseudotypes relative to D614G viruses in both Vero-E6 and 293T-ACE2 cells (**Figure 2A**). As expected for the lower level of ACE2 expression in Vero-E6 cells compared with that in 293T-ACE2 cells (Wang et al., 2021c), the differences in huACE2-Fc sensitivity between D614G and the other viruses were more pronounced in the former cells. No significant differences were observed between the B.1.617.2 and B.1.1.529 pseudotypes in these assays. Using an ELISA, we confirmed that the enhanced sensitivity of B.1.1.529 neutralization relative to that of D614G was due to a higher affinity of huACE2-Fc for the B.1.1.529 spike glycoprotein (**Figure 2B**).

**Figure 2.**
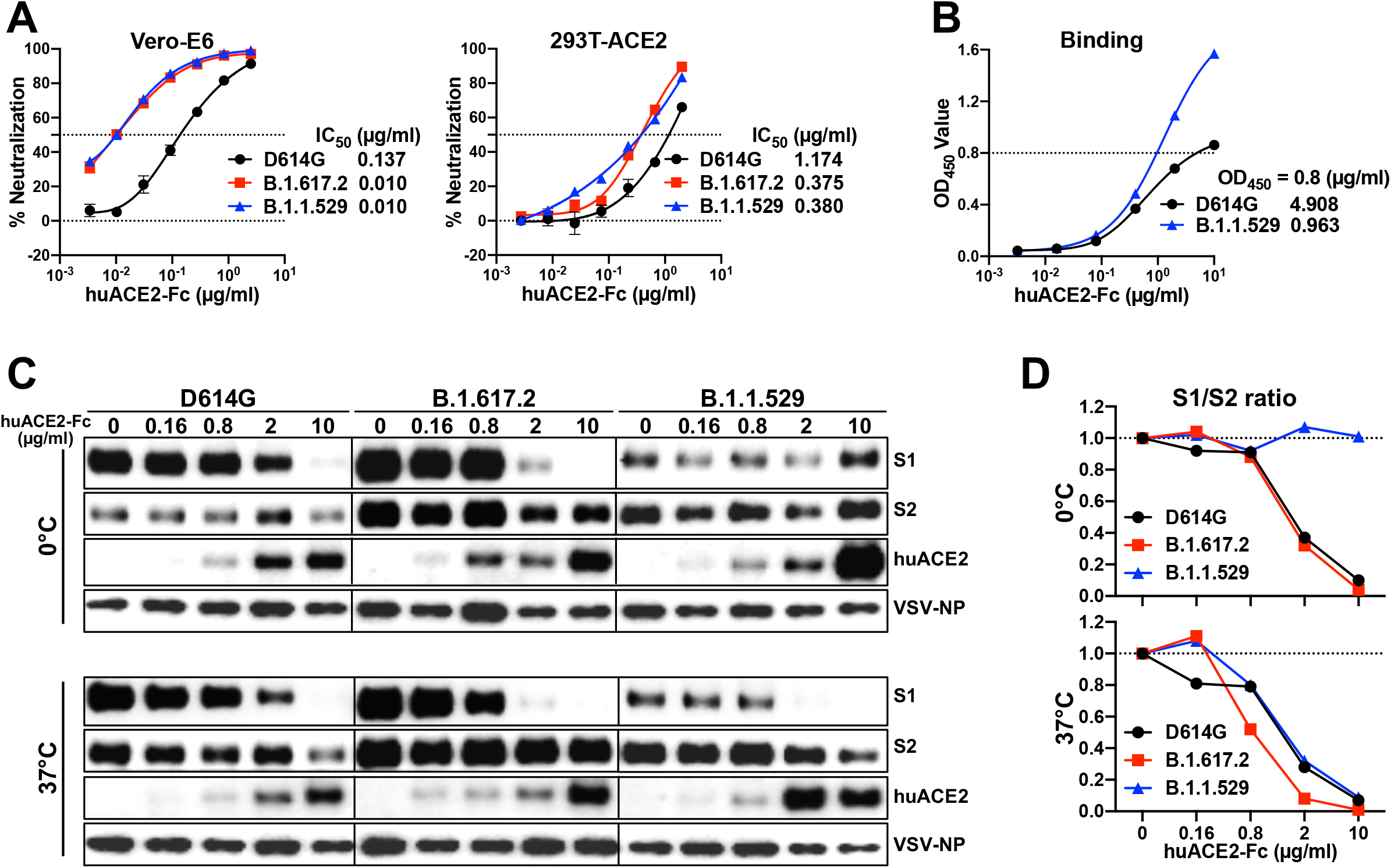
Interaction of soluble ACE2 with the S glycoproteins of D614G, B.1.617.2 and B.1.1.529. **(A)** Neutralization activity of soluble huACE2-Fc against VSV vectors pseudotyped with the indicated S glycoproteins on Vero-E6 (left panel) and 293T-ACE2 (right panel) cells. The results shown in the graphs are the means ± SEM from triplicate measurements and are representative of those obtained in two independent experiments. The huACE2-Fc concentrations resulting in 50% inhibition of infectivity were calculated and are presented to the right of the graphs. **(B)** ELISA binding of soluble huACE2-Fc to the S2P trimers. The data in the graph are the means ± SEM from triplicate measurements obtained in a representative experiment. The concentrations of sACE2-Fc protein required to achieve an OD450 of 0.8 were calculated and are presented to the right of the graph. **(C)** VSV particles pseudotyped with the variant SARS-CoV-2 S glycoproteins were incubated with the indicated concentrations of soluble huACE2-Fc for one hour on ice (upper panel) and 37°C (lower panel). Pelleted VSV particles were analyzed by Western blotting with antibodies against S1, S2, huACE2-Fc, and VSV NP. **(D)** The intensities of the S1 and S2 glycoprotein bands in **(C)** were measured and the S1/S2 ratios for each concentration of sACE2 are shown.

We compared ACE2-induced shedding of S1 from the spike trimers of the D614G, B.1.617.2 and B.1.1.529 pseudoviruses. As differences in soluble ACE2-induced shedding of S1 among some SARS-CoV-2 strains were revealed at 0°C (Wang et al., 2021c), we performed these experiments at 0°C and 37°C. Each variant pseudovirus was incubated with increasing concentrations of huACE2-Fc at either 0°C or 37°C for 1 h and then S1 shedding was quantified by Western blotting (**Figure 2C**). While S1 shedding was similar for all three pseudoviruses at 37°C, the B.1.1.529 spike was significantly more stable at 0°C, with minimal S1 shedding compared to those of the D614G or B.1.617.2 spikes (**Figure 2D**). The observed differences in S1 shedding were not explained by differences in huACE2-Fc binding by the variant S glycoproteins. These results indicate that the ACE2-bound B.1.1.529 S trimer resists the disruptive effects of incubation at 0°C better than the D614G and B. 1.617.2 S trimers.

### Stability of the variant SARS-CoV-2 spikes at different temperatures

Extended periods of incubation at near freezing temperatures can reveal differences in the stability of the spike glycoprotein trimers of SARS-CoV-2 variants (Nguyen et al., 2020; Wang et al., 2021c). We evaluated the effect of temperature on the infectivity of VSV pseudotyped by the D614G, B.1.6172 and B.1.1.529 S glycoproteins. We incubated VSV pseudotypes at 0°C (on ice), 4°C, room temperature (RT), or 37°C for various periods of time before measuring their infectivity on Vero-E6 cells. No differences between the SARS-CoV-2 variants were observed at 4°C, RT, or 37°C; the infectivity of all these variants declined slightly faster at 37°C than at the other temperatures (**Figure 3A**). Notably, at 0°C, the infectivity of the B.1.1.529 pseudotype decayed faster than the infectivities of the D614G or B. 1.617.2 pseudotypes.

**Figure 3.**
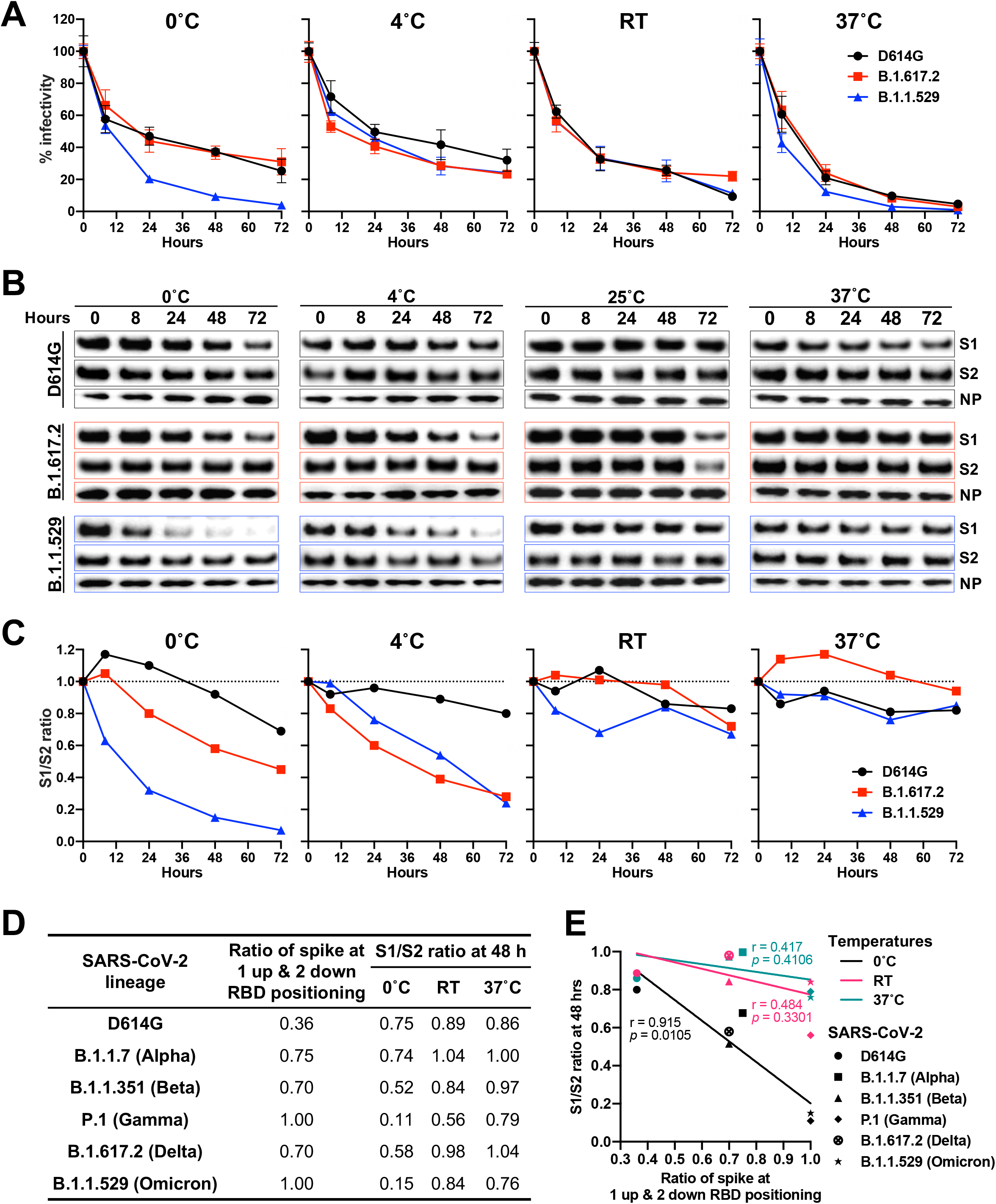
Stability of SARS-CoV-2 D614G, B.1.617.2, and B.1.1.529 spikes at different temperatures. **(A)** VSV vectors pseudotyped by the variant SARS-CoV-2 S glycoproteins were incubated on ice, at room temperature (RT), and at 37°C for various times. The infectivity of the viruses was measured on Vero-E6 cells. The infectivity at each time point is reported, relative to that observed at time 0 for each pseudovirus. Means ± SEM derived from three replicates are shown. The results are representative of those obtained in four independent experiments. **(B)** VSV particles pseudotyped with the variant SARS-CoV-2 S glycoproteins were incubated on ice, at RT, and at 37°C for various times. Pelleted VSV particles were analyzed by Western blotting with antibodies against S1, S2, and VSV NP. The results shown are representative of those obtained in two independent experiments. **(C)** The intensities of the S1 and S2 glycoprotein bands in **(B)** were measured. The S1/S2 ratios are shown for each variant relative to the ratio for pseudoviruses at time 0. **(D)** Relative populations of SARS-CoV-2 variant spikes with one RBD in the “up” position (Cerutti et al., 2021) and S1/S2 ratios after a 48-hour incubation at 0°C, RT and 37°C ((Wang et al., 2021c) and **C** above). **(E)** Correlation between the percentage of spike populations with one RBD in the “up” position and the S1/S2 ratio after a 48-hour incubation at 0°C, RT and 37°C for spike glycoproteins from different SARS-CoV-2 variants. The r and *p* values for each curve are obtained by fitting the data with simple linear regression.

As the observed reduction in infectivity of the B.1.1.529 pseudovirus at 0°C may be due to the shedding of the S1 glycoprotein from the spike, we quantified S1 retention on the viral particles following various periods of incubation at different temperatures (**Figure 3B and C**). At 0°C, a higher rate of spontaneous S1 shedding from the B.1.1.529 pseudoviruses compared with the D614G and B.1.617.2 pseudoviruses was observed, suggesting that spike disassembly may be part of the mechanism for the observed loss of infectivity at this temperature. The integrity of the B.1.617.2 and B.1.1.529 spikes decreased faster than that of D614G at 4°C; as the infectivity of these pseudoviruses differed only modestly at 4°C, mechanisms of spike inactivation that do not result in S1 shedding are likely involved (Wang et al., 2021c).

In a previous study (Wang et al., 2021c), we found that, similar to the B.1.1.529 variant S glycoprotein, the spike glycoprotein of the P.1 (Gamma) SARS-CoV-2 variant also exhibited an unusually high degree of S1 shedding after a two-day incubation on ice. Interestingly, both P.1 and B.1.1.529 variant spike trimers were reported to reside exclusively in a conformational state with only one receptor-binding domain (RBD) in the “up” position (Cerutti et al., 2021; Wang et al., 2021a). We found a correlation between the S1/S2 ratios of different SARS-CoV-2 variant spike trimers at 48 hours of incubation at different temperatures (Wang et al., 2021c) and the population of the variant spike with one RBD in the up position (Cerutti et al., 2021) (**Figure 3D and E**).

### SARS-CoV-2 variants are more resistant to convalescent patient and vaccinee sera

The B.1.1.529 variant has been reported to be significantly more resistant to antibody neutralization than previously characterized SARS-CoV-2 variants (Liu et al., 2021; Planas et al., 2021). We tested the binding of sera from convalescent COVID-19 patients and vaccinees who had received two doses of mRNA-1273 (Moderna) to recombinant D614G and B.1.1.529 spike trimers (S2P) and RBDs (**Figure 4A, 4C, S2, and S3**). Unexpectedly, we observed that while binding to the B.1.1.529 S2P was lower than that to the D614G S2P for all serum samples, the magnitude of the decrease differed significantly between the groups; convalescent patient sera exhibited only a 1.6-fold decrease in binding titer, whereas sera from Moderna vaccinees exhibited a 9.6-fold decrease, on average. We also observed a decrease in binding to the B.1.1.529 RBD compared with that to the D614G RBD for both convalescent patient sera (15.9-fold decrease) and vaccinee sera (44.1-fold decrease); however, the magnitude of the decrease was more similar between the two groups of sera than that seen for binding to the D614G and B.1.1.529 S2P trimers. The vaccinees elicit higher titers of antibodies against the SARS-CoV-2 spike and against the RBD region than the convalescent COVID-19 patients; apparently, the S2P and RBD epitopes targeted by the vaccinee sera are less conserved between the D614G and B.1.1.529 S glycoproteins than the S2P epitopes recognized by the convalescent patient sera.

**Figure 4.**
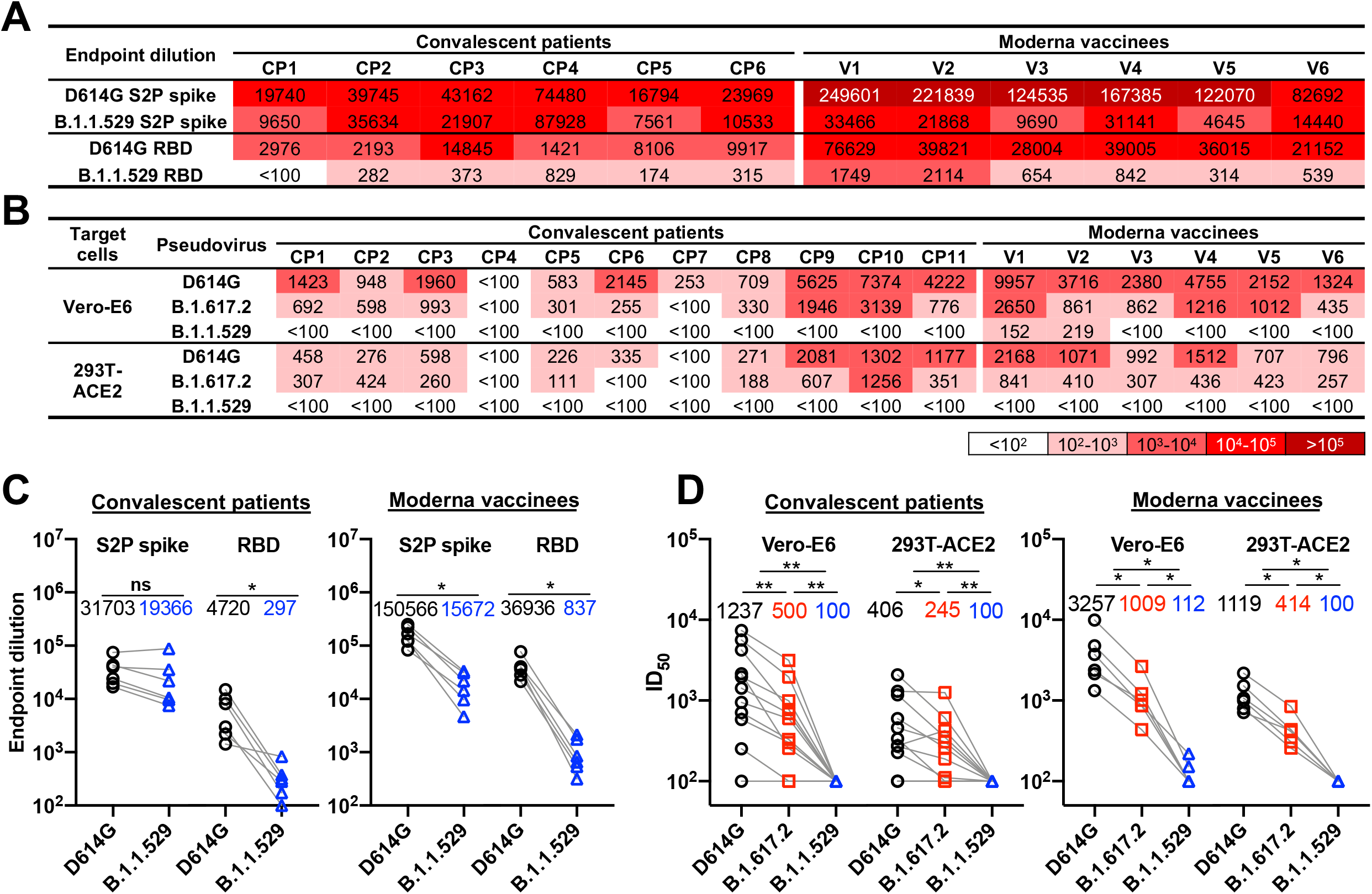
Binding and neutralization sensitivity of SARS-CoV-2 variants. **(A)** The serum binding titers (endpoint dilution) of convalescent COVID-19 patient sera (CP1-CP6) and Moderna vaccinee sera (V1-V6) to the SARS-CoV-2 D614G and B.1.1.529 S2P spike trimers and RBDs. The intensity of shading is proportionate to the titer of binding activity. **See also figure S2 and S3.** **(B)** The serum neutralization titers (ID50) of convalescent patient sera (CP1-CP11) and Moderna vaccinee sera (V1-V6) against VSV particles pseudotyped by the variant SARS-CoV-2 S glycoproteins are reported. The infectivity of the viruses was measured on Vero-E6 cells and 293T-ACE2 cells. The intensity of shading is proportionate to the titer of neutralizing activity. **See also figure S3.** **(C)** Endpoint dilution values of the binding of convalescent patient sera (left) and Moderna vaccinee sera (right) to the B.1.1.529 and D614G S2P trimers and RBDs are compared. **(D)** Serum neutralization ID50 values of convalescent patient sera (left) and Moderna vaccinee sera (right) against VSV particles pseudotyped by B.1.617.2 and B.1.1.529 S glycoproteins were compared with those against the D614G pseudovirus. The geometric means of endpoint dilutions and ID50 values of each group are listed in the graph. Statistical analysis was performed using a paired *t* test (*p < 0.05; **p < 0.01).

To assess the functional consequence of the observed differences in antibody binding to the S glycoprotein trimers, we tested the capability of these sera to neutralize VSV pseudovirus infection of both Vero-E6 and 293T-ACE2 cells (**Figure 4B, 4D, and S4**). The B.1.617.2 pseudovirses were neutralized less effectively than the D614G pseudoviruses by the convalescent patient sera and the vaccinee sera. Neutralization of the B.1.1.529 pseudoviruses by all samples decreased dramatically, with only two samples from Moderna vaccinees having a detectable titer in the Vero-E6 cells. These results indicate that the B.1.1.529 S glycoprotein is significantly less sensitive to neutralization by antibodies elicited to earlier SARS-CoV-2 variant S glycoproteins, either by vaccination or following natural infection.

A comparison of the binding of the convalescent patient and vaccinee sera to the S2P and RBD proteins with the neutralizing activity of these sera suggests differences in the targeted S glycoprotein epitopes. The vaccinees elicit higher titers of antibodies against the SARS-CoV-2 spike and against the RBD region; although this binding correlates better with virus-neutralizing ability, the targeted sites are less conserved between the D614G and B.1.1.529 S glycoproteins. By contrast, many of the S2P epitopes recognized by the convalescent patient sera, although conserved between the D614G and B.1.1.529 S glycoproteins, are not available as neutralization targets on the functional virus spike glycoprotein (Brewer et al., 2021)

## DISCUSSION

The recently emerged B.1.1.529 (Omicron) variant of SARS-CoV-2 has an unusually large number of changes in its spike (S) glycoprotein (**Figure 1A**). Although SARS-CoV-2 components other than S can potentially influence transmission, pathogenesis and immune evasion, the concentration of variation in the B.1.1.529 spike suggests its importance. Indeed, S changes render the virus less sensitive to some therapeutic antibodies and antibodies elicited by current vaccines (Grabowski et al., 2021; Pulliam et al., 2021; Scott et al., 2021). This has raised concerns about the potentially reduced efficacy of vaccines against B.1.1.529 as well as the increased risk for reinfection with this variant (Doria-Rose et al., 2021; Garcia-Beltran et al., 2021; Gruell et al., 2021; Hoffmann et al., 2021; Liu et al., 2021; Planas et al., 2021). Insight into the biology of the novel B.1.1.529 variant would facilitate efforts to manage the COVID-19 pandemic. To this end, we compared the functional properties of the S glycoprotein of B.1.1.529 with that of the globally dominant B.1.617.2 (Delta) variant and the ancestral D614G variant responsible for the early pandemic spread of SARS-CoV-2.

We observed significant differences in the proteolytic processing of the D614G, B.1.617.2 and B.1.1.529 S glycoproteins in expressing cells (**Figure 1B**). Cleavage of coronavirus spike glycoproteins is a prerequisite for activation of their membrane-fusing activity in the infected cells, leading to the formation of short-lived syncytia. The S glycoprotein of the D614G variant, which supplanted the D614 variant responsible for the original SARS-CoV-2 outbreak, exhibited more efficient S glycoprotein cleavage compared with that of its D614 ancestor (Nguyen et al., 2020; Wang et al., 2021c). The S glycoprotein cleavage of most of the SARS-CoV-2 variants of concern is not significantly different from that of the D614 variant (Wang et al., 2021c). Therefore, both the increased efficiency of B.1.617.2 S cleavage and the decreased efficiency of B.1.1.529 S cleavage, relative to that of the D614G S, are noteworthy (**Figure 1B and 1C**). The inefficient processing of the B.1.1.529 S glycoprotein likely contributes to its decreased activity in cell-cell fusion (**Figure 1G**). Syncytium formation represents a major form of the cytopathic effects of SARS-CoV-2, and the decreased cell-fusing capacity of the B.1.1.529 spike glycoprotein could have implications with respect to pathogenesis. Very recent studies indicate that individuals infected with B.1.1.529 are at lower risk of hospitalization and have shorter hospital stays than individuals infected with other SARS-CoV-2 strains, including B.1.617.2 (Mahase, 2021). Although host variables such as age and immune status could potentially influence the observed differences in clinical outcome, intrinsic decreases in B.1.1.529 cytopathicity and pathogenicity may also play a role.

SARS-CoV-2 S glycoproteins incorporated into natural virions and pseudoviruses pass through the Golgi compartment. Relative to the total S glycoproteins synthesized in the producing cell, the virion S glycoproteins are enriched in cleaved glycoproteins modified extensively by complex glycans (**Figure 1D**) (Nguyen et al., 2020; Zhang et al., 2021). This enrichment only partially compensates for the poor processing of the B.1.1.529 S glycoprotein in expressing cells. Nonetheless, despite the lower levels of the B.1.1.529 spike on pseudovirus particles, the infectivity of these pseudoviruses was at least as efficient as those of D614G and B.1.617.2 pseudoviruses (**Figure 1E and 1F**). The ability of the B.1.1.529 S glycoprotein to support virus infection is likely aided by a high affinity for ACE2, a property shared by most emerging SAR-CoV-2 variants of concern (Wang et al., 2021c). Relative to D614G pseudoviruses, both B.1.1.529 and B.1.617.2 pseudoviruses were neutralized more effectively by soluble ACE2 (**Figure 2A**). Direct measurement of the binding of soluble huACE2-Fc to the S glycoprotein trimers of D614G and B.1.1.529 confirmed the higher affinity of the latter spike protein (**Figure 2B**).

The exposure of native SARS-CoV-2 spike glycoproteins to 0°C can reveal strain-dependent differences in the stability and functional integrity of the S glycoprotein trimer. The infectivity of B.1.1.529 pseudoviruses decayed similarly to those of D614G and B.1.617.2 pseudoviruses at room temperature and 37°C, but exhibited a relatively faster decay at 0°C (**Figure 3A**). This corresponded to an increase in the spontaneous shedding of the S1 glycoprotein from the B.1.1.529 spike trimer during prolonged incubation on ice, compared with the D614G and B.1.617.2 spikes (**Figure 3B and 3C**). The cold resistance of enveloped viruses is related to the ability of their oligomeric envelope glycoproteins to resist the destabilizing effects of ice crystal formation (Privalov, 1990). The sensitivity of human immunodeficiency viruses to inactivation on ice is related to the propensity of the viral envelope glycoprotein trimer to undergo conformational transitions from the pretriggered conformation to more “open” receptor-bound conformations (Kassa et al., 2009). Likewise, we found a correlation between cold sensitivity of SARS-CoV-2 variants and the occupancy of the S glycoprotein trimers in a conformation in which one RBD is in the “up” position and the other two RBDs are in the “down” position (**Figure 3D and 3E**). Thus, the cold sensitivity of the B.1.1.529 and P.1 variants may reflect the propensity of the “closed” spike trimer to sample spontaneously the next more “open” state, a conformation with one RBD “up”. This propensity to undergo conformational changes could assist the triggering of conformational transitions in the B.1.1.529 spike and help compensate for a lower spike density on the virion. For the P.1 SARS-CoV-2 variant, which has a more typical spike density, increased triggerability could account for the efficiency with which target cells with low levels of ACE2 can be infected (Wang et al., 2021c).

Relative to the D614G and B.1.617.2 spikes on pseudoviruses, the B.1.1.529 spike demonstrated marked resistance to S1 shedding after exposure to soluble ACE2 at 0°C (**Figure 2C and 2D**). The B.1.1.529 spike shares this property with the P.1 spike but not with the S glycoproteins of other SARS-CoV-2 variants studied (Cerutti et al., 2021; Wang et al., 2021a; Wang et al., 2021c). Apparently, in the more “open” ACE2-bound state, with RBDs in the up position, the B.1.1.529 and P.1 S trimers maintain a tighter association with S1. This property of the B.1.1.529 and P.1 spikes might be a secondary adaptation to an increased spontaneous sampling of the one-RBD-up conformation. During infection of cells, the prolonged stability of the receptor-bound S glycoprotein trimer could allow the requisite number of spike-receptor interactions to be achieved, compensating for a lower density of virion spikes (in the case of B.1.1.529) or lower availability of ACE2 receptors (in the case of P.1). The temperature-dependent regulation of S conformational state could have value for respiratory viruses like SARS-CoV-2, because they infect both cells in the nasal passages that could be at cool temperatures, as well as cells in the lower respiratory tract that are at 37 °C.

The B.1.1.529 pseudoviruses were remarkably resistant to neutralization by sera from convalescent COVID-19 patients and from individuals who had received two doses of the Moderna vaccine (**Figure 4B, 4D and S4**). These results indicate that most of the neutralizing antibodies generated to previous SARS-CoV-2 variants will be less effective against B.1.1.529. Resistant to neutralizing antibodies is clearly a major factor driving the evolution of SARS-CoV-2 variants (Choi et al., 2020). This conclusion is consistent with the increased number of vaccine breakthrough infections involving B.1.1.529 compared with earlier SARS-CoV-2 variants (Andrews et al., 2021; Kuhlmann et al., 2021; Pulliam et al., 2021). The B.1.1.529 spike is apparently well adapted so that any negative consequences of an increased propensity to sample the one-RBD-up conformation, which potentially could result in greater exposure to neutralizing antibodies, are compensated during naturally encountered circumstances.

Although both convalescent patient sera and vaccinee sera failed to neutralize B.1.1.529 pseudoviruses, the binding of the convalescent patient sera to recombinant spike trimer (S2P) of B.1.1.529 was only modestly decreased compared with the binding to D614G S2P. By contrast, the difference in binding of the vaccinee sera to these two recombinant spike trimer was much greater. The magnitude of the decrease in binding to D614G and B.1.1.529 RBDs between the convalescent patient sera and vaccinee sera was not as significant as that seen for binding to the S2P glycoproteins (**Figure 4A and 4C**). The vaccinee sera apparently recognize S2P and RBD epitopes that differ between D614G and B.1.1.529, but which represent targets for neutralization on the functional virus spike. A significant fraction of antibodies in the sera of convalescent patients recognize spike epitopes that are conserved between the D614G and B.1.1.529 S2P glycoproteins but are not available as neutralization targets on the native virus spike. For example, spikes that shed S1 during natural infection could present S2 trimers to the host immune system; many of the antibodies directed against these S2 trimers would fail to bind intact S trimer (Brewer et al., 2021).

In conclusion, we demonstrate that the B.1.1.529 spike glycoprotein has a unique combination of properties differentiating it from previous SARS-CoV-2 variants. In addition to the resistance of the B.1.1.529 spike to antibodies reactive with other SARS-CoV-2 spike glycoproteins, other notable features include the relatively low level of B.1.1.529 S processing and virus spike density, decreased syncytium-forming ability, and the spontaneous and ACE2-regulated stability of the spike trimer, which is temperature-dependent and relates to spike conformational states. A better understanding of the spike-determined biological characteristics of this newly emerged SARS-CoV-2 variant should assist the design of vaccines and other interventions.

## ACKNOWLEDGMENTS

We thank Elizabeth Carpelan for manuscript preparation. This study was supported by a gift from William F. McCarty-Cooper.

## AUTHOR CONTRIBUTIONS

Q.W. and J.G.S. designed the research. Q.W. and S.A. performed the experimental work. S. I., Y.C., L.L. and D.D.H. provided the convalescent and vaccinee sera, and soluble spike trimers. Q.W., S.A., and J.G.S. analyzed the data and wrote the manuscript. All authors reviewed and edited the manuscript.

## DECLARATION OF INTERESTS

The authors declare no competing interests.

## STAR Methods

### EXPERIMENTAL MODEL AND SUBJECT DETAILS

#### Human subjects

Plasma samples were obtained from patients (aged 27-85 years) convalescing from SARS-CoV-2 infection (Wang et al., 2021c). The patients were enrolled in an observational cohort study of patients convalescing from COVID-19 at the Columbia University Irving Medical Center (CUIMC) starting in the spring of 2020. The study protocol was approved by the CUIMC Institutional Review Board (IRB) and all participants provided written informed consent.

Sera were obtained from 6 participants in a phase-I clinical trial of the Moderna SARS-CoV-2 mRNA-1273 vaccine conducted at the NIH, under an NIH IRB-approved protocol (Anderson et al., 2020). The sera used in the present study were obtained after the participants had received two doses of mRNA-1273.

#### Cell lines

HEK293T, 293T-ACE2 (BEI), Vero-E6, and COS-1 cells were cultured in Dulbecco modified Eagle medium (DMEM) supplemented with 10% fetal bovine serum (FBS) and 100 mg/ml penicillin-streptomycin (Thermo Fisher Scientific, Cambridge, MA). Expi293 cells were maintained in suspension culture directly in Expi293TM Expression Medium, supplemented with penicillin and streptomycin, and were incubated at 37°C in a humidified atmosphere of 8% CO2 in air and on a shaker platform rotating at 125 rpm. HEK293T cells, 293T-ACE2 cells, Expi293 cells are of female origin. Vero-E6 and COS-1 are from African green monkey kidneys.

### METHOD DETAILS

#### Plasmid constructs

The codon-optimized SARS-CoV-2 spike (S) gene (Sino Biological, Wayne, PA) encoding the S glycoprotein lacking 18 amino acids at the carboxyl terminus was cloned into the pCMV3 vector. The gene encoding the B.1.617.2 SARS-CoV-2 spike variant was made by introducing additional mutations into the wild-type (D614) S gene using a site-directed mutagenesis kit (Agilent, Santa Clara, CA). The B.1.1.529 S gene was generated by a high-throughput template-guided gene synthesis approach (Liu et al., 2021). The genes encoding the SARS-CoV-2 D614G and B.1.1.529 S trimer ectodomains with 2P and furin cleavage-site changes, and containing a C-terminal 8x His tag (Wrapp et al., 2020) were synthesized and then cloned into the paH vector.

#### Expression and purification of protein reagents

The soluble SARS-CoV-2 S2P spike trimer proteins of the D614G and B.1.1.529 variants were generated by transfecting Expi293 cells with the trimer protein-expressing constructs using FectPRO DNA transfection reagent (Polyplus, New York, NY) and purified from cell supernatants 5 days later using Ni-NTA resin (Invitrogen), according to the manufacturer’s protocol (Liu et al., 2020).

Two μg of each S2P trimer and RBD protein was analyzed on a NuPAGE Bis-Tris protein gel (Invitrogen, Waltham, MA) run at 200 V using MES buffer, after which the gel was stained with Coomassie Blue dye.

#### VSV pseudotyped by SARS-CoV-2 S glycoproteins

To generate VSV-based vectors pseudotyped with SARS-CoV-2 S glycoproteins, 6 X 10^6^ HEK293T cells were plated in a 10-cm dish one day before transfection. Fifteen μg of the SARS-CoV-2 S glycoprotein plasmid was transfected into the HEK293T cells using Polyethylenimine (Polysciences, Warrington, PA). Twenty-four hours later, the cells were infected at a multiplicity of infection of 3 to 5 with rVSV-ΔG pseudovirus bearing a luciferase gene (Kerafast, Boston, MA) for 2 h at 37°C and then washed three times with PBS. Cell supernatants containing pseudoviruses were harvested after another 24 h, clarified by low-speed centrifugation (2000 rpm for 10 min) and filtered through a 0.45-μM filter. Viruses were then aliquoted and stored at −80°C until use (Liu et al., 2020; Wang et al., 2021a; Wang et al., 2021b).

#### Lentiviruses pseudotyped by SARS-CoV-2 S glycoproteins

To generate HIV-based vectors pseudotyped with SARS-CoV-2 S glycoproteins, 6 X 10^6^ HEK293T cells were plated in a 10-cm dish one day before transfection. Then, 7.5 μg of SARS-CoV-2 S glycoprotein expressor plasmid and 7.5 μg pHIV-1_NL4-3_ΔEnv-NanoLuc reporter construct were cotransfected into the HEK293T cells using Polyethylenimine (Polysciences). Cell supernatants containing pseudoviruses were harvested 48 h after transfection, clarified by low-speed centrifugation (2000 rpm for 10 min) and filtered through a 0.45-μM filter. Viruses were then aliquoted and stored at −80°C until use.

#### S glycoprotein expression, processing and incorporation into pseudovirus particles

HEK293T cells were transfected to produce VSV- and HIV-based particles pseudotyped with SARS-CoV-2 S glycoprotein variants, as described above. To prepare viral particles, cell supernatants were collected, centrifuged at low speed (2000 rpm) to remove cell debris, filtered through a 0.45-μM filter and pelleted at 18,000 x g for 1 h at 4°C. In parallel with harvesting the pseudoviruses from the cell supernatants, cells were washed and lysed using 1.5% Cymal-5 at 4°C for 10 min. Cell lysates were then clarified by high-speed centrifugation (18,000 x g for 10 min). Cell lysates and virions were analyzed by Western blotting with the following primary antibodies: rabbit anti-SARS-Spike S1 (Sino Biological, Cat# 40591-T62), rabbit anti-SARS-Spike S2 (Sino Biological, Cat# 40590-T62), rabbit anti-p55/p24/p17 (Abcam, Cambridge, MA (Cat# ab63917)), mouse anti-VSV NP (Millipore, Burlington, MA (Cat# MABF2348)) or mouse anti-GAPDH (Millipore, Cat# CB1001). The Western blots were developed with the following secondary antibodies: HRP-conjugated anti-rabbit antibody (Cytiva, Marlborough, MA (NA934-1ML)) or HRP-conjugated goat antimouse antibody (Jackson ImmunoResearch, West Grove, PA (Cat# 115-035-008)).

S, S1, and S2 band intensities from unsaturated Western blots were calculated using ImageJ Software. S1/S2 ratios represent the ratios of the intensities of the S1 and S2 glycoprotein bands in the Western blots. The values for the processing index of mutant S glycoproteins were calculated as follows:

Processing index = (S1/S × S2/S)_mutant_ ÷ (S1/S × S2/S)_WT_

#### Deglycosylation of S glycoproteins

SARS-CoV-2 S glycoproteins in cell lysates or on pseudovirus particles were prepared as described above. Protein samples were boiled in 1 X denaturing buffer and incubated with PNGase F or Endo Hf (New England Biolabs, Ipswich, MA) for 1 h at 37°C according to the manufacturer’s protocol. The samples were then analyzed by SDS-PAGE and Western blotting as described above.

#### Virus infectivity and stability at different temperatures

The pseudoviruses were freshly prepared as described above, without freezing and thawing. The pseudovirus preparations were incubated with target cells seeded in 96-well plates at a density of 3 X 10^4^ cells/ well. For VSV-based pseudoviruses, Vero-E6 and 293T-ACE2 target cells were cultured for 16-24 hours after infection and then harvested to measure the luciferase activity (Promega, Madison, WI (Cat# E4550)). For HIV-based pseudoviruses, 293T-ACE2 target cells were cultured for 2-3 days after infection and then cells were harvested to measure the NanoLuc luciferase activity (Promega, Cat# N1120).

To measure viral stability at different temperatures, pseudoviruses were incubated on ice, at 4°C, at room temperature and at 37°C for different lengths of time prior to measuring their infectivity in Vero-E6 cells.

To measure the infectivity of pseudovirus variants on target cells expressing different levels of human ACE2, serial dilutions (from 2 μg to 24.7 ng) of the human ACE2 expressor plasmid (Addgene, Watertown, MA (Cat# 1786)) were transfected into 293T cells in 12-well plates using 1 mg/ml PEI. Two days after transfection, cells were trypsinized and used as target cells for measuring the infectivity of VSV pseudotypes (Wang et al., 2021c).

#### Cell-cell fusion assays

For the alpha-complementation assay measuring cell-cell fusion, COS-1 effector cells were plated in black-and-white 96-well plates and then cotransfected with a plasmid expressing alpha-gal and either pCMV3 or the SARS-CoV-1 S glycoprotein variant at a 1:1 ratio, using lipofectamine 3000 (Thermo Fisher) according to the manufacturer’s protocol. At the same time, 293T target cells in 6-well plates were cotransfected with plasmids expressing omega-gal and human ACE2 at a 1:1 ratio, using lipofectamine 3000. Forty-eight hours after transfection, target cells were scraped and resuspended in medium. Medium was removed from the effector cells, and target cells were then added to effector cells (one target-cell well provides sufficient cells for 50 effector-cell wells). Plates were spun at 500 x g for 3 min and then incubated at 37 °C in 5% CO2 for 4 h. Medium was aspirated and cells were lysed in Tropix lysis buffer (Thermo Fisher Scientific). The β-galactosidase activity in the cell lysates was measured using the Galacto-Star Reaction Buffer Diluent with Galacto-Star Substrate (Thermo Fisher Scientific), following the manufacturer’s protocol.

#### ELISA assays

Fifty nanograms of S2P spike trimer or RBD protein (Sino Biological Inc. (Cat: 40592-V08H121 and 40592-V08H)) was coated on each well of an ELISA plate at 4°C overnight. Then the plates were blocked with 300 mL of blocking buffer (1% BSA and 10% bovine calf serum in PBS) at 37°C for 2 h. Afterwards, serially diluted huACE2-Fc (Sino Biological Inc. (cat# 10108-H02H)), convalescent serum, or Moderna vaccinee serum was added and incubated at 37°C for 1 h. Next, 100 mL per well of 10,000-fold diluted Peroxidase AffiniPure goat anti-human IgG (H+L) antibody (Jackson ImmunoResearch) was added and incubated for 1 h at 37°C. Between each step, the plates were washed with PBST three times. Finally, the TMB substrate (Sigma, St. Louis, MO) was added and incubated for 5 min at room temperature before the reaction was stopped using 1 M sulfuric acid. Absorbance was measured at 450 nm.

The concentrations of huACE2-Fc that achieved an OD450 of 0.8 and and the serum endopoint dilution that achieved an OD450 value > 3-fold over background were calculated by fitting the data in five-parameter dose-response curves in GraphPad Prism 9 (GraphPad Software Inc., San Diego, CA).

#### S1 shedding from spike trimers

VSV particles pseudotyped with S glycoprotein variants were prepared as described above. To evaluate spontaneous S1 shedding at different temperatures, the cell supernatants containing pseudoviruses were incubated on ice, at 4°C, room temperature and 37°C for different times. Virus particles were then pelleted at 18,000 x g for 1 h at 4°C. The pelleted samples were resuspended in 1 X LDS sample buffer (Invitrogen, Cat# NP0008) and analyzed by Western blotting.

To evaluate soluble huACE2-Fc-induced S1 shedding, the cell supernatants containing pseudovirus particles were incubated with soluble huACE2-Fc (Sino Biological Inc (cat# 10108-H02H)) at different concentrations on ice and 37°C for 1 h. Afterwards, virus particles were pelleted at 18,000 x g for 1 h at 4°C. The pelleted virus particles were washed once with cold PBS before the samples were resuspended in 1 X LDS sample buffer and analyzed by Western blotting. S1, S2 and NP were detected as described above; huACE2-Fc bound to the pseudovirus particles was detected with Peroxidase AffiniPure goat anti-human IgG (H+L) antibody (Jackson ImmunoResearch).

#### Neutralization assay

Sera were collected from March to June 2020 from New York City patients that recovered from COVID-19; six sera were collected from Moderna vaccinees (Wang et al., 2021b). Pseudovirus neutralization assays were performed by incubating VSV vectors pseudotyped by S glycoprotein variants with serial dilutions of huACE2-Fc (cat# 10108-H02H), sera from convalescent COVID-19 patients or vaccinee sera in triplicate in 96-well plates for 1 h at 37°C. Approximately 3 x 10^4^ target cells (Vero-E6 or 293T-ACE2 cells) per well were then added. The cultures were maintained for an additional 16-24 h at 37°C before luciferase activity was measured as described above. Neutralization activity was calculated from the reduction in luciferase activity compared with mock-treated controls. The concentrations of huACE2-Fc and serum titers that inhibit 50% of infection (the IC50 and ID50 values, respectively) were determined by fitting the data in five-parameter dose-response curves in GraphPad Prism 9 (GraphPad Software Inc., San Diego, CA).

## QUANTIFICATION AND STATISTICAL ANALYSIS

Evaluations of statistical significance were performed employing Student’s unpaired or paired *t* test using GraphPad Prism 9 software. Levels of significance are indicated as follows: ns, *p*>0.05; \**p*<0.05; \*\**p*<0.01; \*\*\**p*<0.001; and \*\*\*\**p*<0.0001. Linear correlation was determined by fitting the data with simple linear regression. EC50, IC50 and ID50 values were determined by fitting the data in five-parameter dose-response curves in GraphPad Prism 9. Western blot data were analyzed by Image Lab and ImageJ software. All data presented is representative or mean data derived from at least two independent experiments.

## SUPPLEMENTAL INFORMATION TITLES AND LEGENDS

**Figure S1.** The infectivity of VSV vectors pseudotyped by the variant SARS-CoV-2 S glycoproteins on 293T cells transfected with the indicated amounts of an ACE2-expressing plasmid. Four independent experiments are shown and data are shown as the means and standard deviations. **See also figure 1E**.

**Figure S2. SDS-polyacrylamide gel of purified D614G and B.1.1.529 S2P spike trimers. See also figure 2B, figure 4A and figure 4C**.

**Figure S3. Binding of convalescent COVID-19 patient sera (CP1-CP6) and Moderna vaccinee sera (V1-V6) to the SARS-CoV-2 D614G and B.1.1.529 S2P trimers (A) and RBD proteins (B).** The S2P trimers and RBD proteins were captured on ELISA plates. Serially diluted sera (starting from 1:100) were incubated with the plates and the bound antibodies were measured. Dashed lines show a level 3-fold above the background OD450 values. Data are presented as means ± SEM. **See also figure 4A and 4C**.

**Figure S4. Neutralization of VSV pseudotypes by sera from convalescent COVID-19 patients and vaccinees.** Neutralization curves are shown of convalescent COVID-19 patient sera (CP1-CP11) and Moderna vaccinee sera (V1-V6) against VSV particles pseudotyped by the variant SARS-CoV-2 S glycoproteins on Vero-E6 **(A)** and 293T-ACE2 **(B)** cells. Data are represented as means ± SEM. **See also figure 4B and 4D.**

## Notes

### Competing Interest Statement

The authors have declared no competing interest.

