## Supplemental figures for "Functional properties of the spike glycoprotein of the emerging SARS-CoV-2 variant B.1.1.529"

### Figure S1

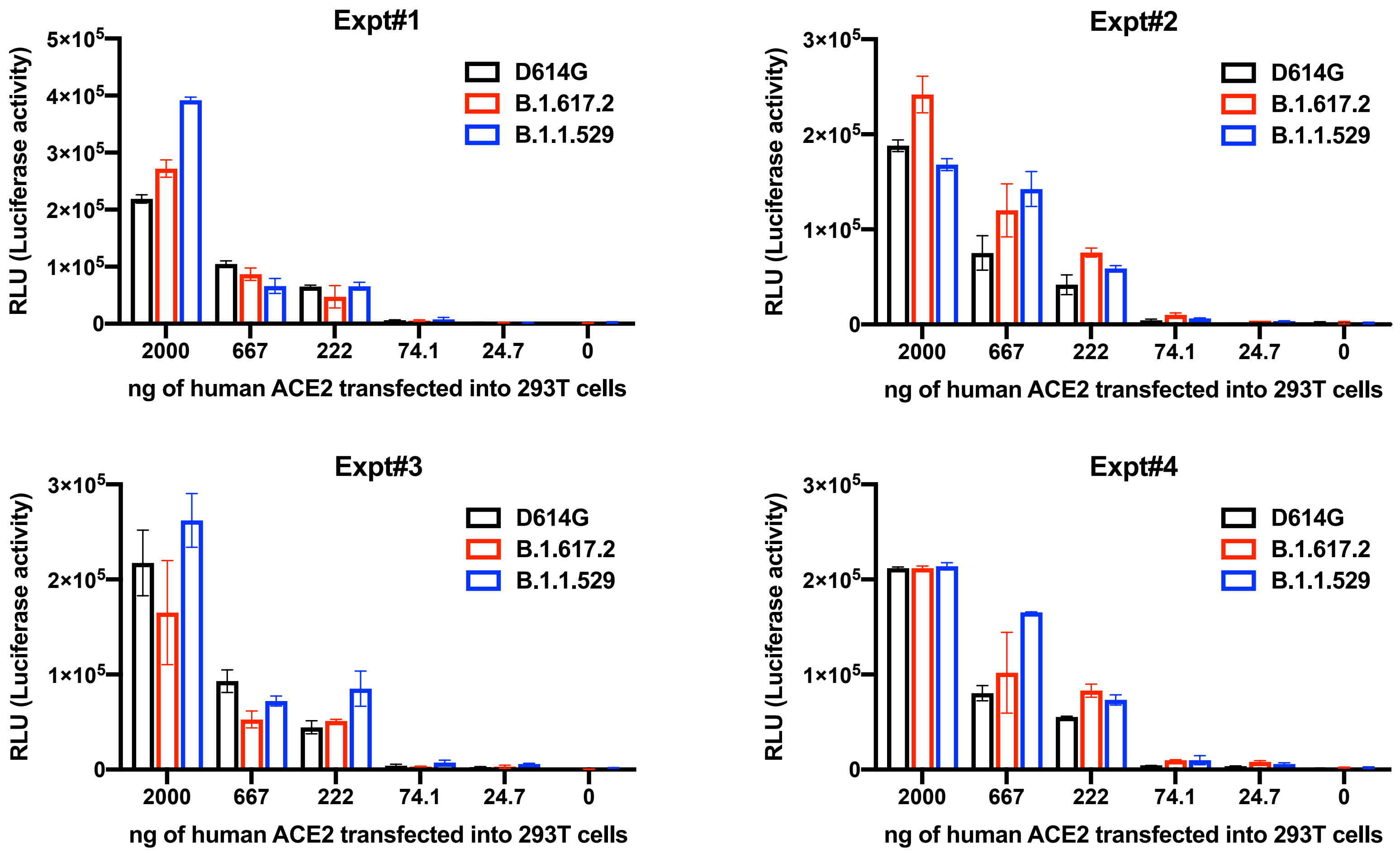


**Figure S1.** The infectivity of VSV vectors pseudotyped by the variant SARS-CoV-2 S glycoproteins on 293T cells transfected with the indicated amounts of an ACE2-expressing plasmid. Four independent experiments are shown and data are shown as the means and standard deviations. **See also figure 1E**.

### Figure S2

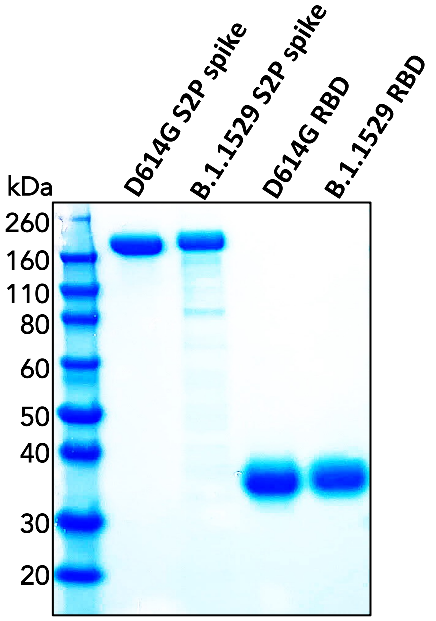


**Figure S2. SDS-polyacrylamide gel of purified D614G and B.1.1.529 S2P spike trimers. See also figure 2B, figure 4A and figure 4C**.

#### Figure S3

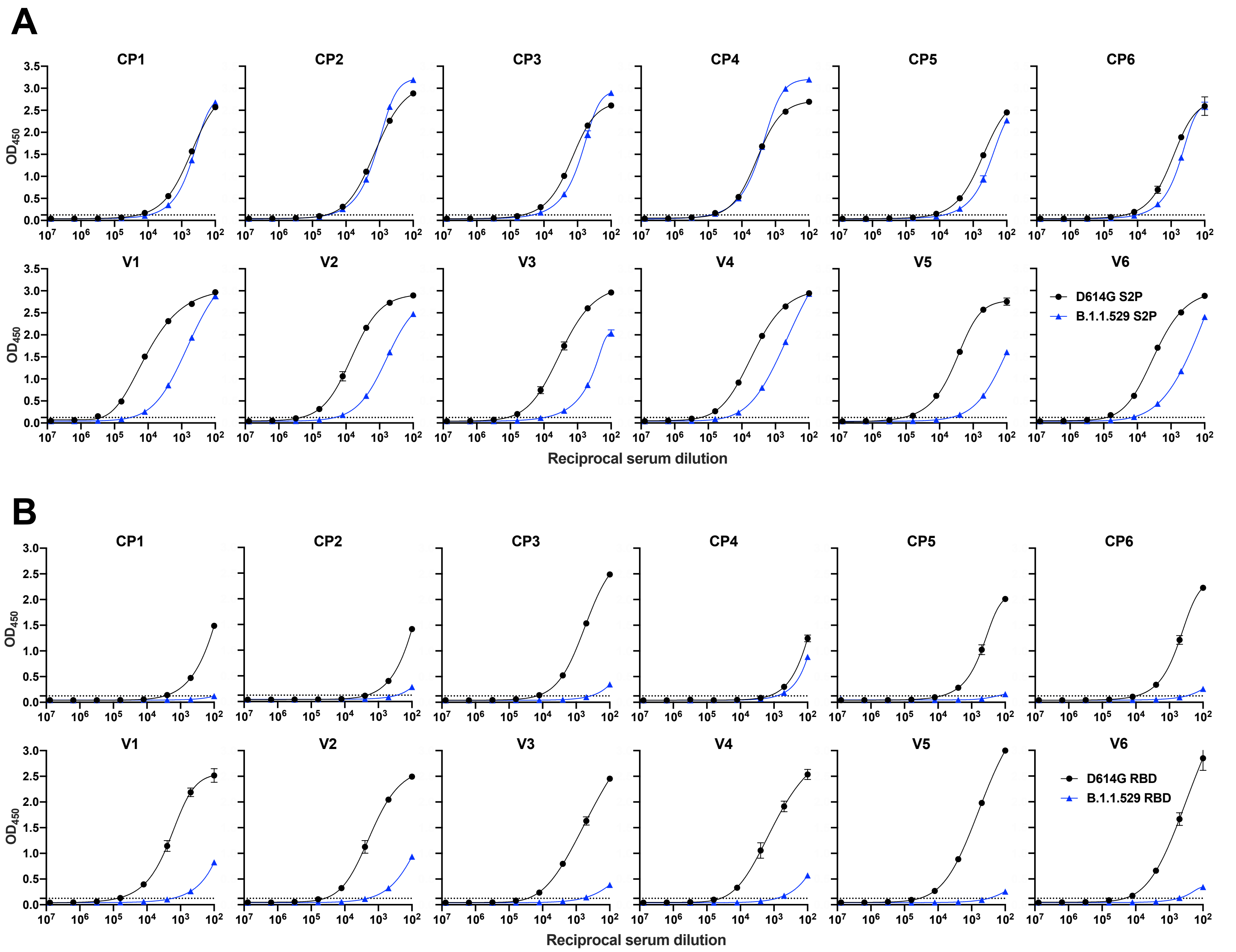


**Figure S3. Binding of convalescent COVID-19 patient sera (CP1-CP6) and Moderna vaccinee sera (V1-V6) to the SARS-CoV-2 D614G and B.1.1.529 S2P trimers (A) and RBD proteins (B).** The S2P trimers and RBD proteins were captured on ELISA plates. Serially diluted sera (starting from 1:100) were incubated with the plates and the bound antibodies were measured. Dashed lines show a level 3-fold above the background $OD_{450}$ values. Data are presented as means ± SEM. **See also figure 4A and 4C**.

### Figure S4

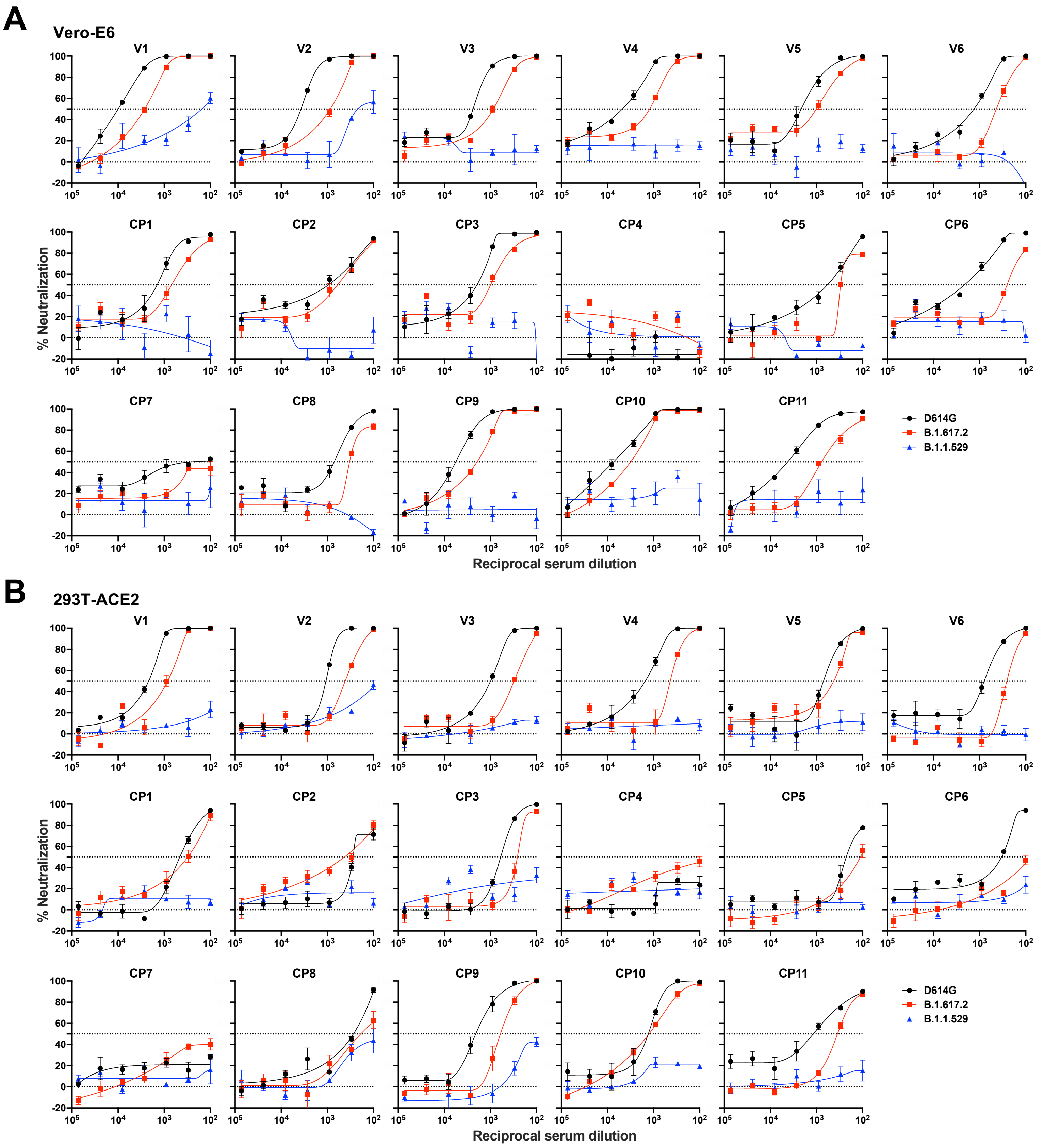


**Figure S4. Neutralization of VSV pseudotypes by sera from convalescent COVID-19 patients and vaccinees.** Neutralization curves are shown of convalescent COVID-19 patient sera (CP1-CP11) and Moderna vaccinee sera (V1-V6) against VSV particles pseudotyped by the variant SARS-CoV-2 S glycoproteins on Vero-E6 **(A)** and 293T-ACE2 **(B)** cells. Data are represented as means ± SEM. **See also figure 4B and 4D.**
